# Control of eNOS and nNOS through regulation of obligatory conformational changes in the reductase catalytic cycle

**DOI:** 10.1101/410571

**Authors:** John C. Salerno, Benjamin L. Hopper, Dipak. K. Ghosh, Israel M. Scott, Jonathan L. McMurry

## Abstract

Endothelial and neuronal nitric oxide synthases (eNOS, nNOS) are important signal generators in a number of processes including angiogenesis and neurotransmission. The homologous inducible isoform (iNOS) occupies a multitude of conformational states in a catalytic cycle, including subnanosecond input and output states and a distribution of ‘open’ conformations with average lifetimes of ~4.3 ns. In this study, fluorescence lifetime spectroscopy was used to probe conformational states of purified eNOS and nNOS in the presence of chaotropes, calmodulin, NADP^+^ and NADPH. Two-domain FMN/oxygenase constructs of nNOS were also examined with respect to calmodulin effects. Optical biosensing was used to analyze calmodulin binding in the presence of NADP^+^ and NADPH. Calmodulin binding induced a shift of the population away from the input and to the open and output states of NOS. NADP^+^ shifted the population towards the input state. The oxygenase domain, lacking the input state, provided a measure of calmodulin-induced open-output transitions. A mechanism for regulation by calmodulin and an elucidation of the catalytic mechanism are suggested by a ‘conformational lockdown’ model. Calmodulin speeds transitions between input and open and between open and output states, effectively reducing the conformational manifold, speeding catalysis. Conformational control of catalysis involves reorientation of the FMN binding domain, of which fluorescence lifetime is an indicator. The approach described herein is a new tool for biophysical and structural analysis of NOS enzymes, regulatory events and other homologous reductase-containing enzymes.

**A note to the reader:** This manuscript has over the past several years been submitted to and rejected by several journals, usually on the basis of reviewer opinion that it was not an important enough result to merit inclusion in the journal. Owing to the passing of the first author and the loss of his expertise in fluorescence lifetime spectroscopy, it has become too onerous a task to continually revise the manuscript to suit the whims of reviewers who nevertheless still reject the work. We are thus simply releasing the final form of the manuscript to BioRxiv in the hopes that it finds a readership who will find, as we do, that the results are of value to the field.

## Introduction

Synthesis of the signaling molecule nitric oxide by the endothelial and neuronal nitric oxide synthase isoforms (eNOS and nNOS) requires the delivery of three electrons from NADPH via a three domain reductase unit that comprises the C terminal portion of the enzymes. The endothelial isoform functions as a signal generator in cardiovascular system homeostasis [1, 2], insulin secretion, in the control of cardiac function, angiogenesis, and other processes, while the NO produced by nNOS acts as a neurotransmitter in the central nervous system [3-5], and as a signal in skeletal muscle. Calmodulin (CaM) regulates NO synthesis by eNOS and nNOS; the stimulation of electron transfer through flavin cofactors to the heme catalytic site is mediated by the intracellular Ca^2+^ influx [6].

The reductase unit of eukaryotic NOS is homologous to P450 reductase and includes NADPH, flavin adenine dinucleotide (FAD) and flavin mononucleotide (FMN) binding domains [6-14]. A canonical CaM binding site is located between the oxygenase domain and the FMN binding domain. Both eNOS and nNOS have an approximately 42 residue autoinhibitory element (AI) as an insertion in the FMN binding domain and an extended C-terminal element that restricts electron transfer [15-18]. In contrast, the inducible isoform (iNOS) produces NO as a cytotoxin in immune response [19-21] and is synthesized in response to cytokines. Lacking the AI and the extended C terminal inhibitory region, iNOS binds CaM very strongly and is not Ca^2+^ regulated [22].

CaM controls NO synthesis in eNOS and nNOS by regulating electron transfer. A formal tethered shuttle mechanism was proposed in which the FMN binding domain dissociates from a reductase complex ‘input state’ and reorients to transfer electrons to the oxygenase domain [7, 23, 24]. Work from other groups also supports shuttle mechanisms in which reductase function requires major conformational changes, e.g., [12, 25-31]. In addition to switching on the production of nitric oxide from a negligible rate, CaM binding to nNOS holoenzyme [27, 32-35] increases NADPH cytochrome c reductase activity, typically by a factor of four, and significantly increases steady state flavin fluorescence [7, 12, 32-35].

iNOS holoenzyme and truncated constructs thereof exist in solution in multiple conformational states that can be resolved by FMN fluorescence lifetimes [36]. An input state that is likely the solution counterpart of homologous crystal structures was identified, with FMN and FAD forming a closely coupled chromaphoric dimer with a 90 ps fluorescence lifetime. An output state in which FMN is quenched by interaction with ferriheme can be detected in the truncated two-domain iNOS oxyFMN construct with a lifetime of 0.9 ps. The low quantum yield of this state is the reason for the low steady state fluorescence of FMN in iNOS holoenzyme. A heterogeneous distribution of ‘open’ states in which the FMN binding domain is associated with neither the two domain (FAD/NADPH) dehydrogenase unit nor the oxygenase domain can be detected with an average lifetime of ~4.3 ns. The kinetics cycle of the reductase unit requires the FMN binding domain to traverse states of this type as it moves between the input and output configurations.

Truncated NOS oxyFMN constructs (containing only the FMN and heme domains) have been useful in examining FMN domain-oxygenase interactions without interference from interactions between the FMN binding domain and the two domain dehydrogenase unit consisting of the NADPH and FAD binding domains, which dominate in holoenzyme [23-27, 37]. The absence of the low quantum yield FAD-FMN dimer of holoenzyme makes oxyFMN more fluorescent than holoenzyme [36, 38]. Quenching of FMN fluorescence by ferriheme provides a means to probe interdomain interactions in NOSoxyFMN. These observations demonstrate conclusively that FAD-FMN interactions produce a ~90 ps state, that free or open configuration FMN domain had a majority 4.3 ns state, and that FMN-heme interactions produce a ~0.9 ns state. This does not imply that these are the sole fluorescence states; it appears that there are minority intermediate lifetime states of FMN that do not require heme. FAD fluorescence contributes to the 90 ps state and perhaps to shorter lived states not observable with our instruments.

CaM activation of nNOS produces a shift in the conformational distribution that reduces the population of the 90 ps input state and increases population of the open and output states [39]. These observations suggest that CaM releases the FMN binding domain from the reductase complex, favoring longer-lived (and hence higher quantum yield) output and open states.

This study describes experiments with bovine eNOS and rat nNOS holoenzymes and oxyFMN constructs. FMN fluorescence provides a description of obligatory conformational changes associated with the catalytic cycle, and allows examination of the effects of activation on different steps of the conformational cycle. Obligatory conformational intermediates can be resolved using appropriate constructs and logically assigning species with different lifetimes and their correlation to NOS catalytic cycles. The approach described herein should allow advances in the knowledge of catalysis and control of NOS enzymes by providing information about important conformational states, opening new avenues for biophysical and structural analysis of this important enzyme, and in addition advance understanding of other enzymes containing homologous reductase catalytic units.

## Materials and Methods

cDNA encoding rat nNOS holoenzyme, a gift from Dr. S. Snyder (Johns Hopkins, MD), was cloned into pCWori+ [40, 41]. Rat nNOS was expressed in *E. coli* strain BL21DE3 and purified using ammonium sulfate precipitation followed by 2’, 5’-ADP Sepharose chromatography or nickel chromatography [39, 41, 42]. Activity was measured by oxyhemoglobin assay, and was 500-700 nmols/min/mg protein [41-43]. Purified nNOS contained 0.8-1.0 heme/mol (CO difference spectrum extinction coefficient of 74 mM^-1^ cm^-1^), and FMN and FAD contents after extraction from nNOS were at least 90% of heme. Rat nNOSoxyFMN was expressed and purified as reported earlier [42]. It consists of the oxygenase and FMN binding domains, and is truncated directly after the negatively charged tail of the final alpha helix of this domain. OxyFMN constructs lack the FAD and NADPH binding domains [42]. Bovine eNOS was similarly expressed in *E. coli* strain BL21DE3, and purified using 2’, 5’-ADP Sepharose chromatography [44].

Rapid kinetic experiments were performed using an Applied Photophysics SX stopped flow unit. Reactions were initiated by mixing 4 μM solutions of nNOS, 1 mM arginine, and 200 μM NADPH in air saturated bis-tris propane (BTP) with 10% glycerol, 100 mM NaCl at pH 7.5 with 120 μM calmodulin and 1 mM CaCl_2_. Kinetics of steady state fluorescence changes in response to calmodulin were performed by adding 10 μM calmodulin and 0.5 mM CaCl_2_ to 1 μM nNOS in 50 mM MOPS, pH 7.4, 50 mM NaCl by simultaneously injecting both solutions into a cuvette; kinetics in these experiments were on the time scale of seconds and minutes. Similar slow kinetic experiments were performed by mixing with NADP^+^ and excess EDTA.

Excitation and emission spectra were recorded on a PTI (now Horiba Scientific) Quantamaster spectrophotometer at 23° C. Excitation and emission slits were both set at 5 nm, and a PMT voltage of 600V was used. Samples were measured in a 1.2 ml quartz cuvette with a path length of 1 cm. The fluorescence of constructs was measured at a concentration of 2 μM in 40 mM BTP containing 1 mM DTT (pH 7.4). Flavin fluorescence emission spectra were measured by exciting samples at 473 nm, and fluorescence intensity were measured from 480 nm to 680 nm. All spectra were corrected for instrumental artifacts by subtracting the baseline emission spectrum of the buffer.

Time-resolved intensity decays were recorded using a PicoQuant Fluotime 100 or a PTI Picoquant time-correlated single-photon counting (TCSPC) fluorescence lifetime spectrometer as described earlier [36, 39, 45]. The excitation at ~ 440 nm was obtained using a pulsed laser diode with 20 MHz repetition rate; experiments conducted with excitation at 378 nm and 473 nm with similar lasers produced similar results. The excitation was vertically polarized and the emission was recorded through a polarizer oriented at 54.7°, the magic angle. Appropriate long-pass filters from Chroma Technology Group (Rockingham, VT) were used to eliminate scattered excitation light.

The fluorescence intensity decays were analyzed as the sum of individual single exponential decays:

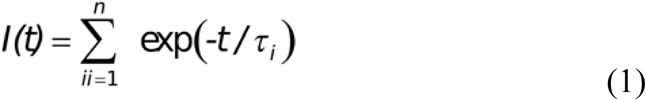

where the *τ*_*i*_ are the decay times and *α*_*i*_ are the amplitudes. The fractional contribution of each component to the steady-state intensity is:

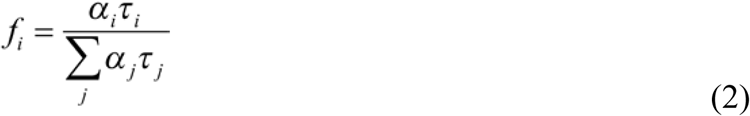

The mean (intensity weighted) lifetime is represented by:

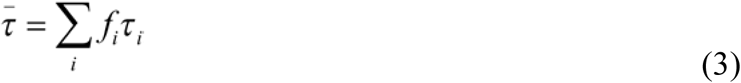

and the amplitude weighted lifetime is given by:

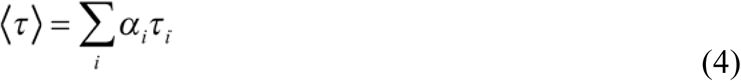

The values of *α*_*i*_ and *τ*_*i*_ were determined using the PicoQuant Fluofit 4.1 (Professional Version) software with the deconvolution of instrument response function and nonlinear least squares fitting. The goodness-of-fit criterion was determined by the χ^2^ value. See also [46].

Biolayer interferometry was performed using a ForteBio Octet QK and streptavidin sensors as described [47].

## Results

Steady state FMN fluorescence in NOS is increased by CaM binding or addition of chaotropes that weaken protein-protein interactions [26, 38, 48, 49]. This is widely recognized as an indicator of conformational changes associated with CaM activation. Fluorescence emission spectra of FMN in all three NOS holoenzymes and FMN containing constructs are dominated by a broad band with a peak around 525 nm. The steady state fluorescence intensity of NOS holoenzymes is weak (less than 20%) in comparison to oxyFMN constructs or independently expressed FMN binding domains (based on total FMN); our work on iNOS showed that this is caused by formation of a short lived, low quantum yield FAD-FMN chromaphoric dimer in the majority state of the holoenzyme [36]. In iNOS and iNOS-derived constructs, FMN fluorescence can be represented as the sum of an electron input state with a lifetime of 90 ps because of strong FAD-FMN coupling, a series of open states in which FMN does not interact strongly with other cofactors and has a flavodoxin-like fluorescence lifetime of 4.3 ns, and an output state in which quenching by heme produces a lifetime of 0.9 ns. We previously reported that activation of nNOS by Ca^2+^/CaM favors the open and output states at the expense of the input state [39]. The nNOS states are similar to those observed in iNOS, except that it was possible to observe the ~1 ns component in nNOS holoenzyme. Resolution of this component in iNOS required the use of truncated constructs.

The short lifetimes of the input and output states reflect quenching of FMN through interactions with FAD and heme, respectively. Energy transfer to other groups quenches steady state fluorescence by shortening the lifetime; the Förster equation describing the rate of exciton transfer in s^-1^ via dipole-dipole interaction is

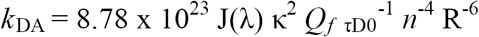

where *Q*_f_ is the fluorescence quantum yield of the isolated donor, κ^2^ =(**a**^.^**d**-3(**a**^.^**r)(d**^.^**r**))^2^ is the dipole orientation factor, _τD0_ is the isolated acceptor lifetime, *n* is the refractive index of the surroundings, J(λ) is the spectral overlap integral, and R is the effective donor-acceptor distance in Å. The spectral overlap integral for FMN and FAD is ~ 4.6 x 10^-15^ cm^3^ M^-1^ [50]; the overlap integral for FMN and high spin heme is ~ 0.9 x 10^-13^ cm^3^ M^-1^, because the FMN emission spectrum overlaps the ferriheme and bands.

The radiative lifetime of FMN, calculated from absorbance and emission spectra, is 15-18 ns [50, 51]; the quantum yield of free FMN, with a lifetime of 4-5 ns, is therefore 20-30%. Small flavoproteins with no strong quenchers are similar. The isolated NOS FMN binding domain has a majority component with a lifetime of 4.3 ns and, like the open states of holoenzyme, has quantum yield of ~25%. The 1 ns output state has a quantum yield of ~ 6%, and the 100 ps input state has a quantum yield of less than 1%. The input state corresponds to the reductase crystalfstructures of nNOS and P450 reductase in which FAD and FMN are in Van der Waals contact [52, 53]. This state might be better described as a chromophoric dimer than by using the Förster equation.

Figure 1A shows the temperature dependence of FMN fluorescence decay in nNOS. As the temperature is lowered from 37° C to 4° C, the population of the 90 ps state decreases and the 0.9 ns and 4.3 ns state populations increase. This suggests that the entropy of the input state is significantly lower than the entropies of the open state and output state, consistent with the exposure of a larger surface area in these states because of the breakup of the input state complex, requiring the formation of a larger shell of ordered water.

**Figure 1.**
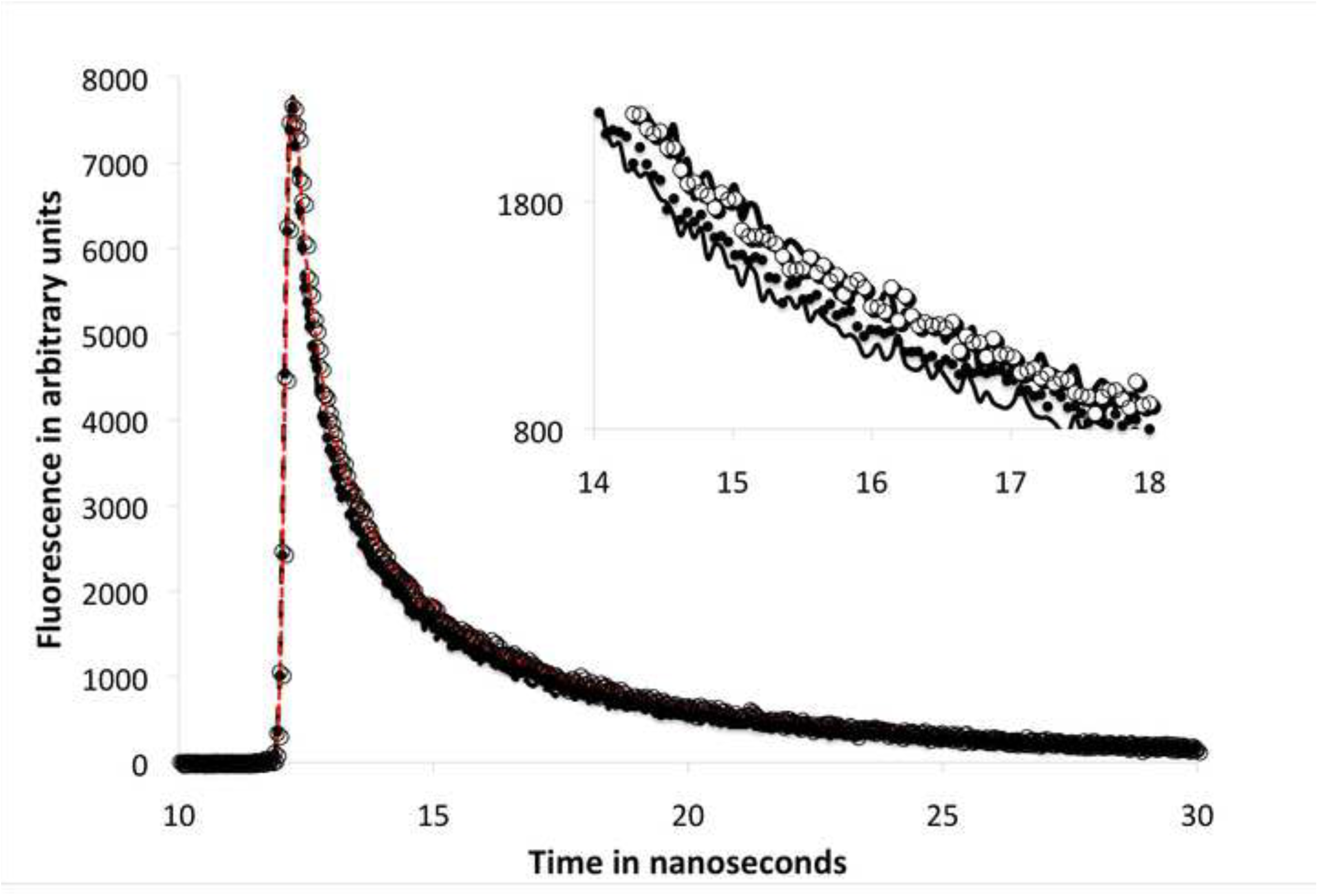
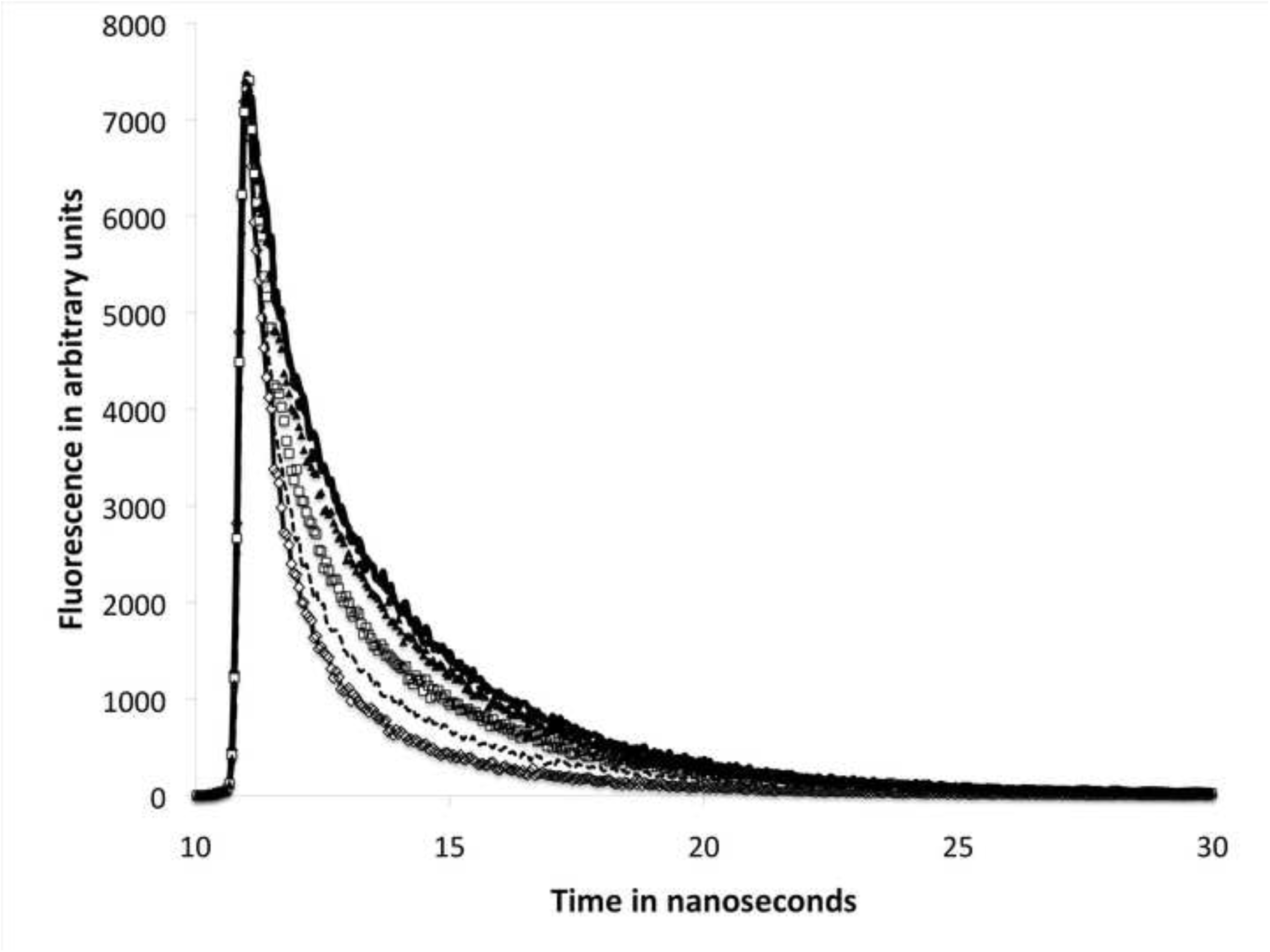
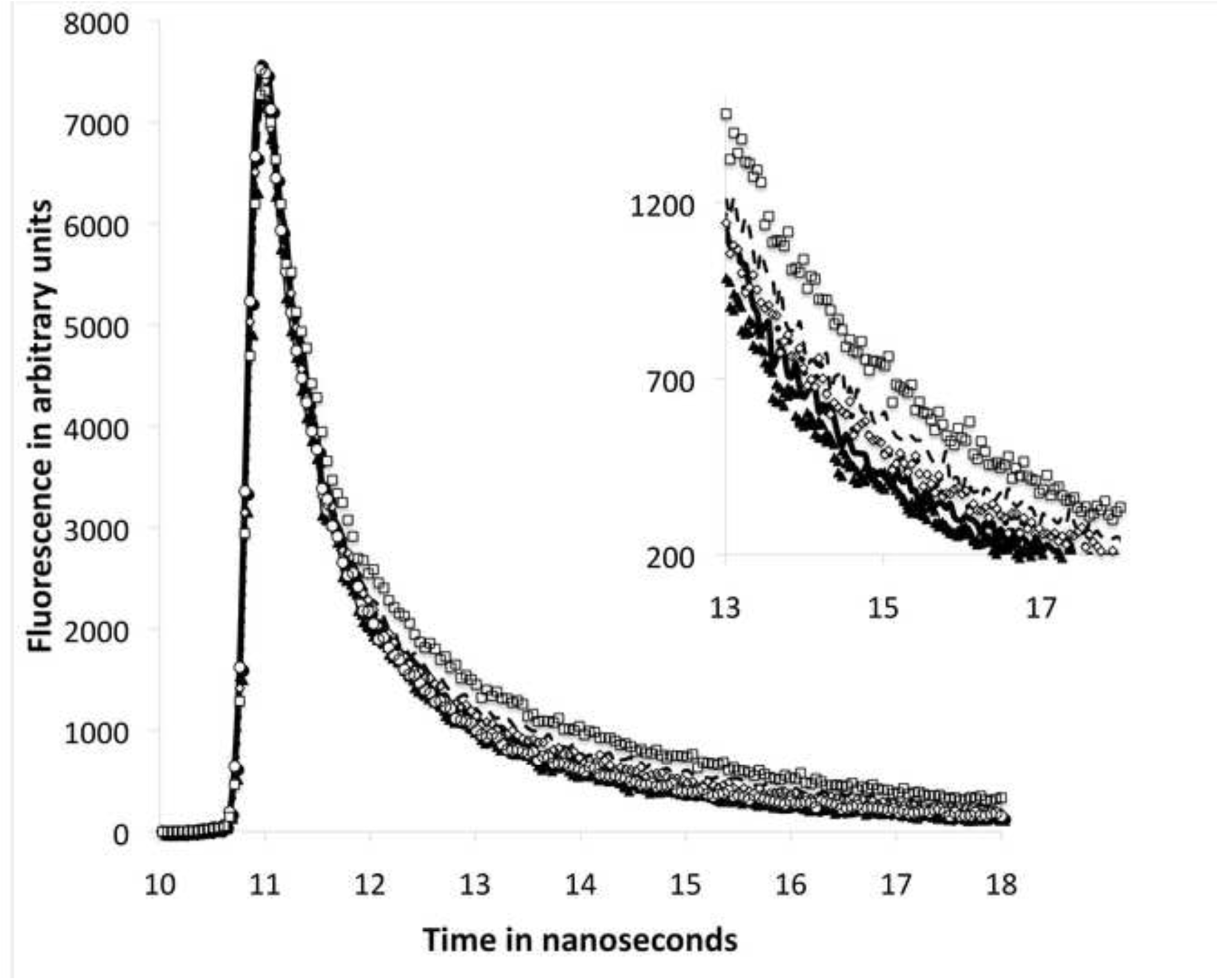
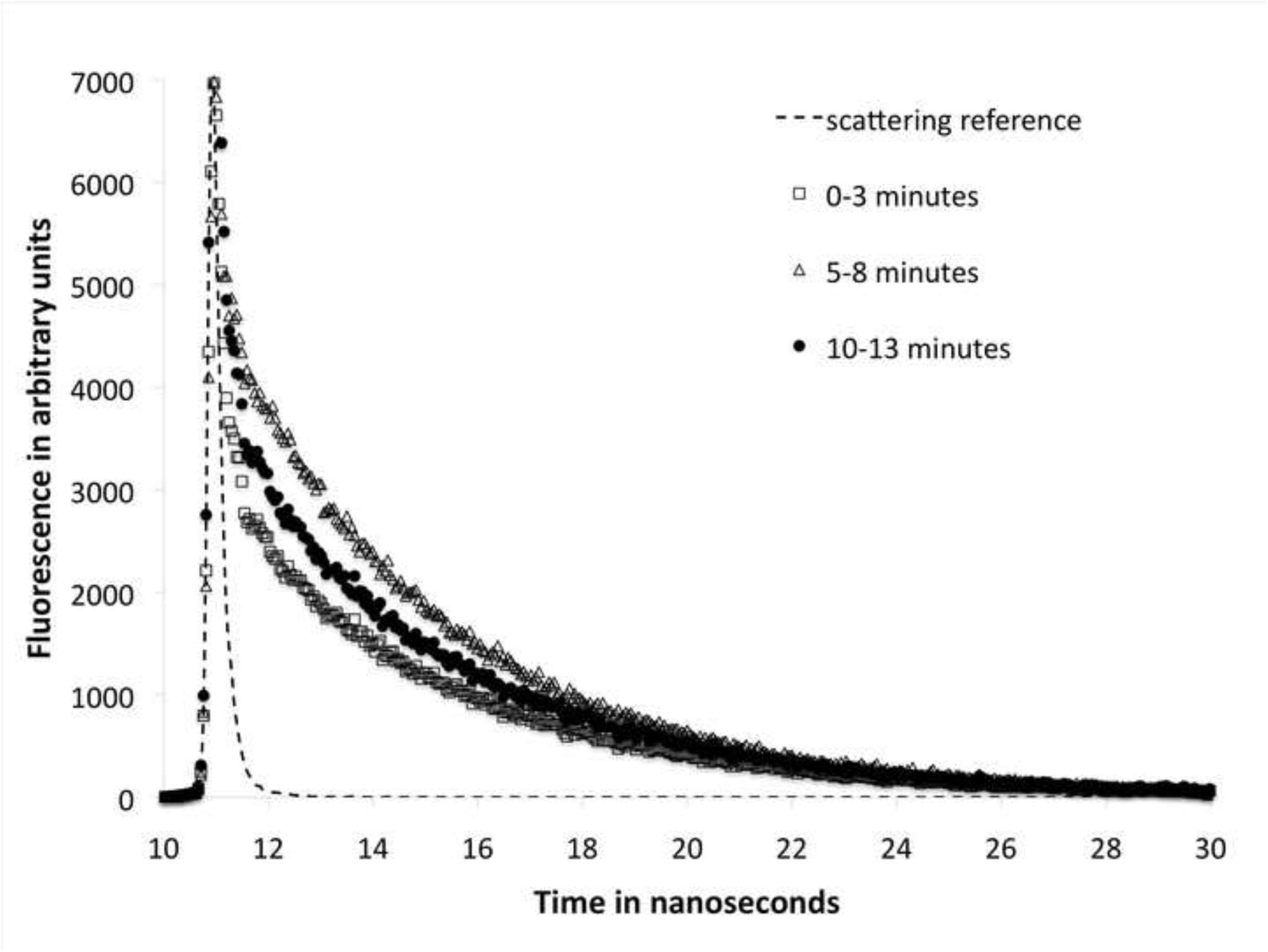
(A) Temperature dependence of FMN fluorescence decay in nNOS showing the decrease in the population of the 90 ps state as the temperature is lowered from 37° C to 4° C. Decays were obtained for 2 µM nNOS in 50 mM MOPS, pH 7.4, 50 mM KCl, 10% glycerol. –, 37°C; •, 25°C; Ο, 10°C; **—,** 4°C. (B) Effects of chaotropes on the fluorescence decays of nNOS. Addition of perchlorate progressively lead to FMN domain release in the 0.5 M to 4 M concentration regime. ◊ nNOS as in (A); - - - 1M sodium perchlorate; ◻ 1.6 M perchlorate; ▴ 2 M perchlorate; • 3 M perchlorate; – 4 M perchlorate. (C) Urea effects on 2 µM nNOS. Urea concentrations shown are ▴0.25 M, ◯ 0.5 M, ◊ 5M, – 8M, and ◻ 12 M. (D) Unpacking effects on 2 µM eNOS holoenzyme at 25°C. Holoenzyme was diluted into sample buffer and decays were collected at the times shown. It takes approximately 3 minutes to collect each trace.

Figure 1B shows the effects of chaotropes on the fluorescence decays of nNOS. Chaotropes have been shown to increase the FMN and tryptophan fluorescence of NOS and to increase the cytochrome c reductase activity, but they are not able to promote significant levels of NO synthesis. As shown here, perchlorate produces significant increases in the long-lived species associated with open conformations in which FMN is exposed to solvent and not closely associated with heme or FAD. Guanidine has similar effects (data not shown). Urea is much less effective (Figure 1C), but produces some increases in longer lived states at high concentrations. NaCl and KCl at molar concentrations had no effect. The primary effects of chaotropes on NOS arise from destabilization of the short-lived input state, resulting in increased population of long-lived, high quantum yield open states. NO synthesis is not supported because the output state cannot be reached.

Highly concentrated nNOS and eNOS samples are much more stable than dilute enzyme. When such samples are diluted from 50-100 μM to 1-2 μM for spectroscopy, they are initially largely in the input state, and it requires at least 10 minutes at 4° C to approach a steady state conformational distribution in the absence of calmodulin. An example of this ‘unpacking’ effect is shown in Fig. 1D. Unpacking is faster in the presence of Ca^2+^/CaM. We attribute this effect, and the enhanced stability of the enzyme, to the formation of aggregates in which the NOS reductase domains interact, stabilizing the input state.

Figure 2 shows fluorescence decays for eNOS holoenzyme. The holoenzyme has components that correspond closely to those associated with nNOS. As in nNOS, FMN fluorescence emission at 525 nm decays as a multiple exponential, and reasonable fits require at least three components. Lifetimes are again approximately 90 ps, 0.9 ns, and 4.3 ns, with the majority 90 ps state typically accounting for just over half the population. The largest fractional occupation we observed in eNOS was just over 60%, considerably less than typical nNOS preparations. The long tail again appears heterogeneous.

**Figure 2.**
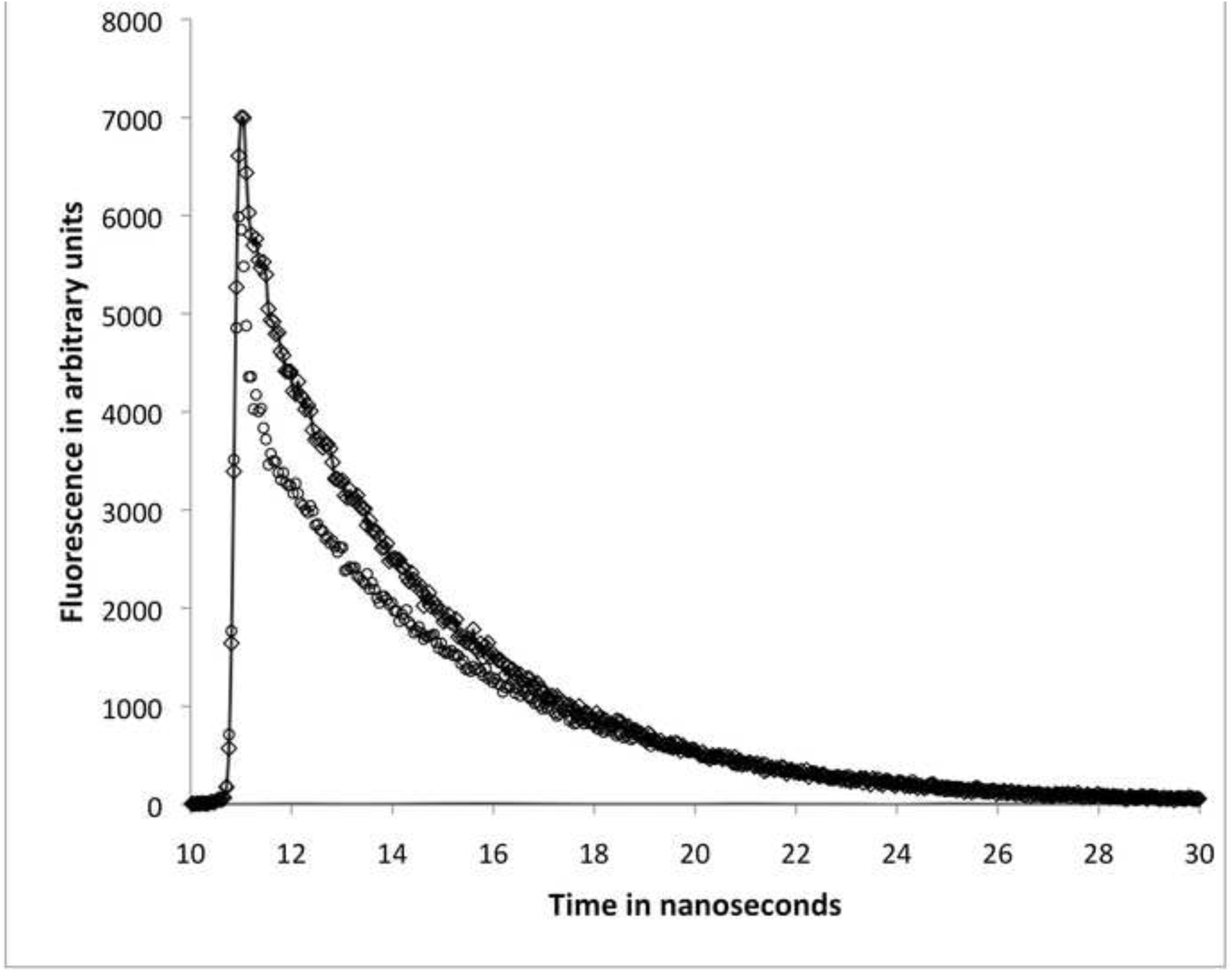
Fluorescence decays for eNOS holoenzyme showing the effect of Ca^2+^/CaM addition. Sample was prepared as in Figure 1, except that 2 μM eNOS was used. The top trace contained 10 μM CaM and 100 μM CaCl_2_.

As in nNOS, CaM binding decreases the fractional population of the 90 ps input state significantly, with a concomitant increase in the population of the 0.9 ns output state. Addition of CaM to Ca^2+^-depleted eNOS produces a partial result, likely due to endogenous Ca^2+^ in our CaM preparations. Addition of Ca^2+^ increases the effect of CaM.

Interestingly, addition of EDTA to CaM-activated nNOS and eNOS lowered activity without reversing the effect of Ca^2+^/CaM on the conformational distribution. It is possible to repeatedly activate and deactivate a NOS preparation by alternatively adding Ca^2+^ and EDTA. EDTA rapidly releases CaM from eNOS and nNOS, but this is not accompanied by a rapid return to the original conformational distribution. One reason for this is that the effect on the rates of transition between states is primary. EDTA-induced release of CaM from nNOS immediately halts equilibration within the manifold, freezing in the conformational distribution of the activated enzyme. Fluorescence decays slowly, approaching the inactivated level in about thirty minutes.

The time frame of conformational effects associated with activation is explored in Figure 3. Figure 3A shows stopped flow absorbance results obtained by mixing 4 μM nNOS, arginine and NADPH with 120 μM CaM and 1 mM Ca^2+^_;_ the high CaM concentration is necessary to obtain rapid, diffusion limited binding. The rapid phase of absorbance changes measured as A_428_ - A_415_ corresponds to formation of the ferrous oxy heme compound of nNOS at the expense of ferriheme. Additional changes at longer times are associated with the accumulation of the ferrous NO complex. Oxygen binding is very rapid compared with reduction, which here is rate limiting. The dotted line is a single exponential with a rate constant of 23 s^-1^, corresponding to a half time of 30 ms. This is in reasonable agreement with more extensive modeling of nNOS turnover through several cycles using a system of differential equations in a previous communication; rates of heme reduction were 40-50 s^-1^ [10]. These results indicate that activation via CaM binding takes place within 10-20 ms of mixing, consistent with the diffusion limited rate of CaM binding, and the output state forms within 50 ms.

**Figure 3.**
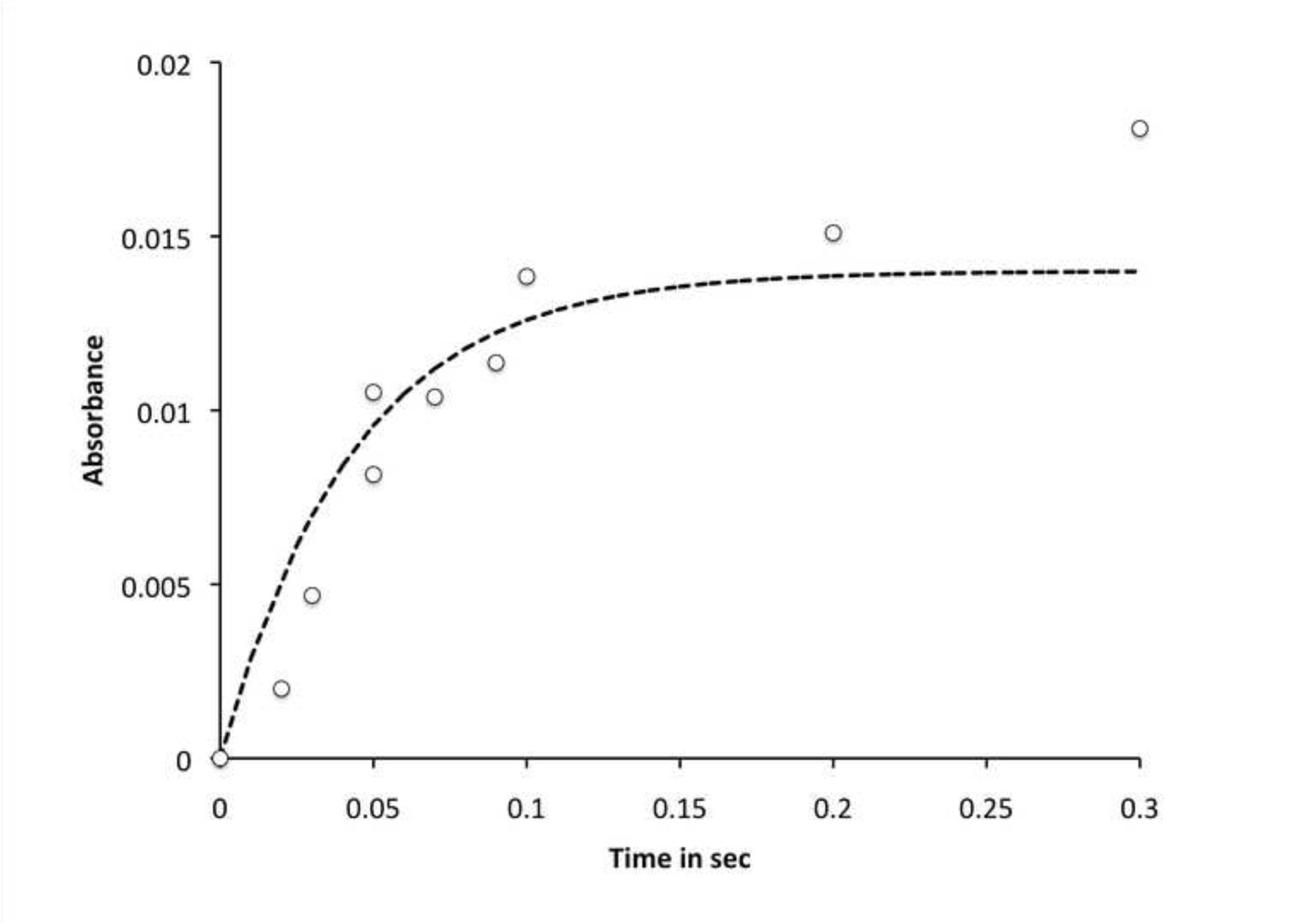
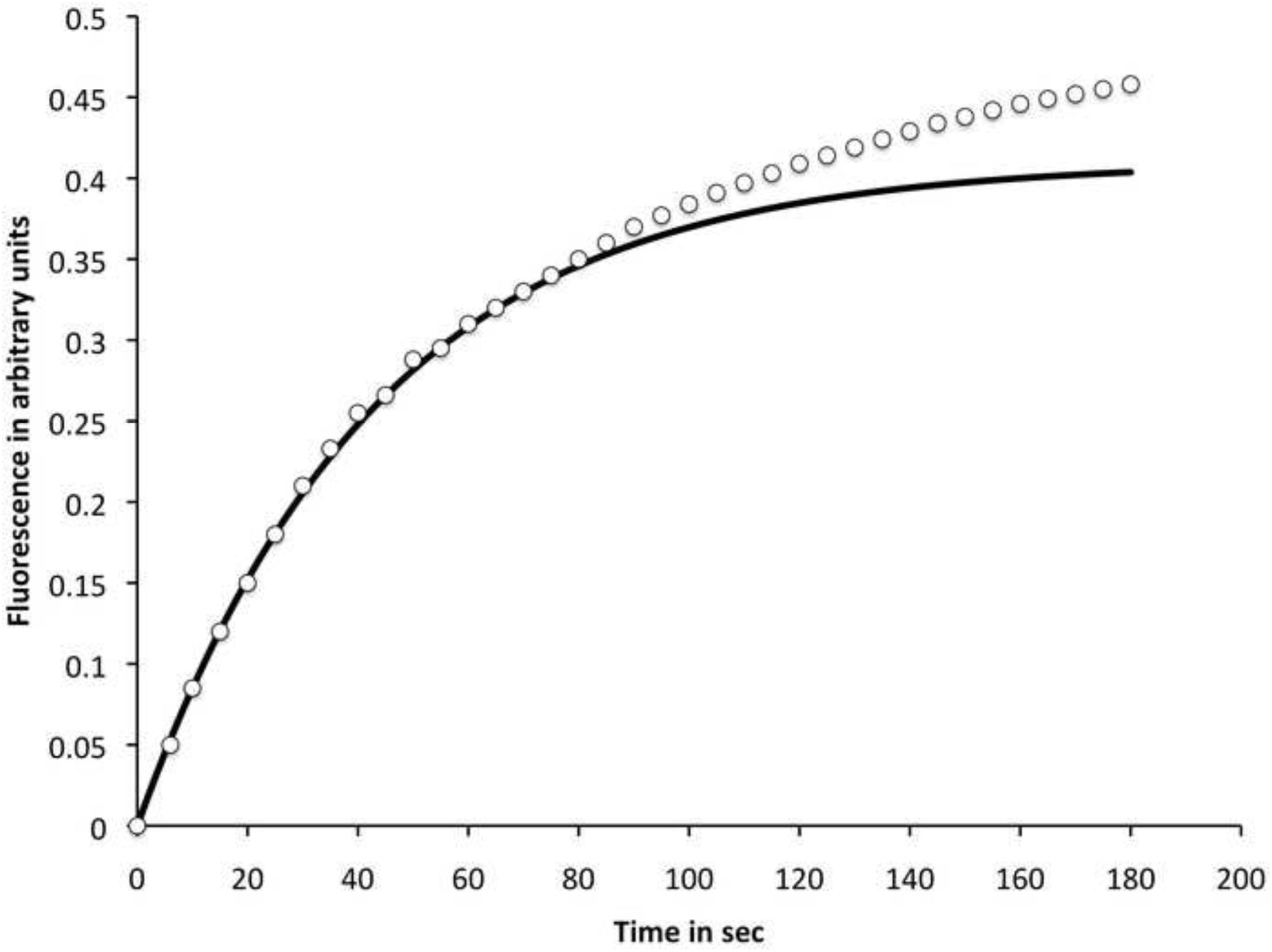
Time frame of conformational effects associated with activation. (A) stopped flow absorbance results (426 nm) obtained by mixing 4 μM nNOS, arginine and NADPH with 120 μM CaM and 1 mM Ca^2+^. The dotted line is a single exponential with a rate constant of 23 s^-1^. (B) Time course of steady state fluorescence changes in nNOS after activation with 20 μM CaM and 0.5 mM CaCl_2_; excitation wavelength 450 nm, emission wavelength 525 nm. The solid line is an exponential with a half time of 30 s.

In contrast, Figure 3B shows the time course of steady state fluorescence changes after CaM activation. Although under these conditions catalysis is initiated in less than a second, the development of additional fluorescence occurs with a half time of 30 s (exponential line). There are additional slower components. The clear implication is that the steady state fluorescence changes reported previously do not correspond to activation by CaM. Instead, they occur gradually after activation as a result of it. This somewhat counter-intuitive finding is the result of the domination of steady state fluorescence by long-lived open conformations, and will be discussed later.

The results for the nNOS two domain oxyFMN construct are shown in Figure 4. Elimination of the 90 ps input state by truncation of the FAD and NADPH binding domains produces a long lifetime (~4.3 ns) majority state, very similar to the long lifetime state in the holoenzyme. In the trace shown the FMN decays as a pure 4.5 ns component; a fit line is completely obscured by the data points. Addition of CaM increases the proportion of the shorter lifetime (1 ns) output state; here the slightly more rapid decay of fluorescence can be accounted for by assigning 10% of the enzyme to the output state. This accounts for the opposite effect of CaM on steady state fluorescence of nNOSoxyFMN and holoenzyme; the increase in holoenzyme fluorescence is due to the shift away from the short lived, weakly fluorescing input state, which is not present in oxyFMN construct because it lacks an FAD binding domain. The oxyFMN construct is much more fluorescent than holoenzyme as a result, and CaM binding reduces its fluorescence.

**Figure 4.**
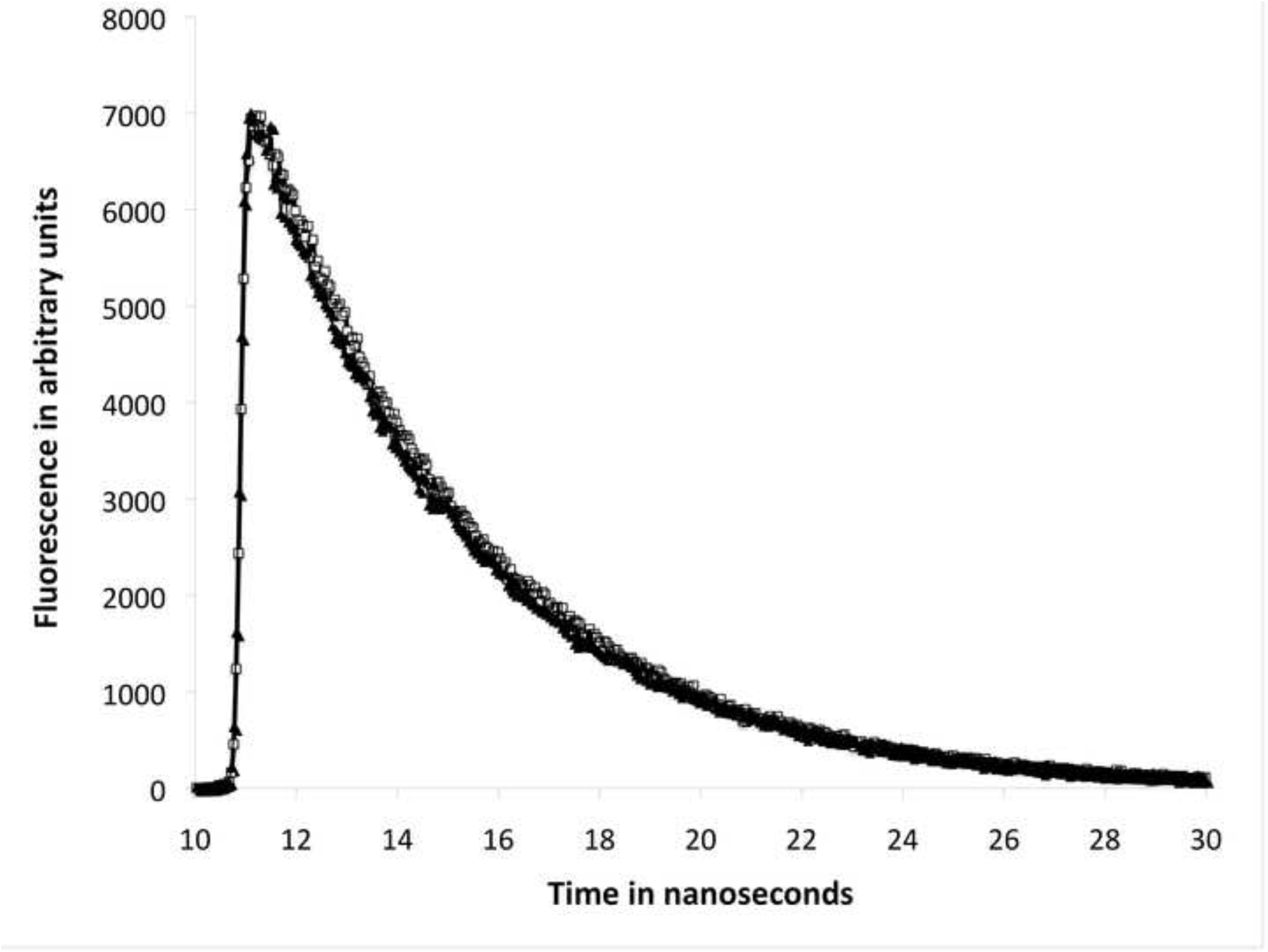
Fluorescence decays for the nNOS two domain oxyFMN construct showing the effect of Ca^2+^/CaM addition. Conditions as in Figure 1 except that the sample contained 2 μM nNOS oxyFMN construct at 25°C. ◻, oxyFMN; ▴, oxyFMN + 4 μM CaM and 10 μM Ca^2+^.

We note that in iNOSoxyFMN coexpressed with CaM, 30% of the construct was in the output state. A significantly larger fraction of nNOSoxyFMN was observed in the output state in other experiments (at least 20% by electron paramagnetic resonance, EPR), and it is possible that in other preparations of the construct a larger yield of the 0.9 ns state could be obtained. It is not possible to do fluorescence and EPR experiments under the same conditions; both the redox state of the chromophores and the concentration of enzyme must be different.

The majority open state FMN fluorescence in oxyFMN is similar to highly fluorescent flavoproteins, and these long-lived states dominate steady state fluorescence even in holoenzyme because of their high quantum yield, roughly 50 times that of the input state. The flavin isoalloxazines in the input state crystal structures are closely associated, consistent with the short observed lifetimes.

As previously reported, the FMN fluorescence lifetimes of independently expressed nNOS FMN domain are similar to flavodoxin. The FMN domain has a majority state with a lifetime of 4.3 ns; a minority component has a lifetime of ~ 2 ns. This corresponds well to the long lifetime states of NOS holoenzymes and oxyFMN constructs, and confirms our assignment of these states to ‘free’ FMN binding domains in the open conformation.

Table 1 summarizes the data for lifetime experiments with eNOS and nNOS constructs and holoproteins. Essentially similar results were obtained with excitation wavelengths that pumped all three major isoalloxazine bands; data shown were obtained with 440 nm excitation. Similar results have been obtained using 375 nm and 478 nm excitation. In all cases intensity decays were fitted with a multi-exponential function. Parameters for three component fits are shown; in all experiments at least two well-resolved components are observed.

**Table 1.** - Lifetimes and component amplitudes of FMN fluorescence in eNOS and nNOS.

| Sample | A<sub>1</sub> | $\tau_1$<br>(ns) | A<sub>2</sub> | $\tau_2$<br>(ns) | A<sub>3</sub> | $\tau_3$<br>(ns) | $\chi^2$ |
| --- | --- | --- | --- | --- | --- | --- | --- |
| nNOS-full length [39] | 0.38 | 4.3 | 0.19 | 0.9 | 0.43 | 0.09 | 1.12 |
| nNOS-full length (immediately after dilution into MOPS buffer) | 0.16 | 4.3 | 0.03 | 0.9 | 0.82 | 0.09 | 0.99 |
| nNOS-full length+CaM [39] | 0.53 | 4.3 | 0.33 | 0.9 | 0.14 | 0.09 | 1.31 |
| nNOS-HoloProtein G810 [39] | 0.7 | 4.1 | 0.3 | 0.72 |  |  | 1.21 |
| nNOS-HoloProtein G810 + CaM [39] | 0.65 | 4.2 | 0.35 | 0.48 |  |  | 1.12 |
| nNOS-oxyFMN | 1 | 4.5 |  |  |  |  | 1.01 |
| nNOS-oxyFMN + CaM | 0.9 | 4.4 | 0.1 | 0.9 |  |  | 1.11 |
| eNOS-full length | 0.36 | 4.2 | 0.1 | 0.9 | 0.54 | 0.09 | 1.08 |
| eNOS-full length + CAM | 0.63 | 4.2 | 0.21 | 0.9 | 0.16 | 0.09 | 1.24 |

Amplitudes (“A_x_”) are the fractional populations of the states. Experimental traces are highly reproducible, but errors arising from fitting uncertainties reduce the confidence in both the lifetimes and amplitudes to about 10%; in the case of minor components the uncertainty is larger (e.g.: A_2_ for eNOS in the best fit was 0.10, but nearly as good fits could be obtained with 20% larger or smaller components. A_2_ for the nNOS full length immediately after dilution best fit was 0.03, but nearly as good fits could be obtained with values between 0 and 0.05. Variability in preparations is addressed in the text. Some values are from a prior report [39]and are noted.

The majority components for eNOS and nNOS holoenzyme have lifetimes of 90 ps, similar to what we recently reported for iNOS holoenzyme, and like iNOS the eNOS and nNOS holoenzymes also have a long lifetime (4.3 ns) state assigned to open conformations. This component accounts for majority of the steady state intensity, but only about 10-15% of the enzyme population in eNOS and nNOS. Unlike iNOS holoenzyme, a significant intermediate lifetime component can be detected in both eNOS and nNOS holoenzymes after activation; this component represents a significant fraction of FMN in the oxyFMN construct of both iNOS and nNOS, but is at most 5% in iNOS holoenzyme. Because of severe proteolysis, we cannot present reliable results for eNOS oxyFMN.

Preparations of both eNOS and nNOS exhibit significant variability in the populations of the states observed here. In nNOS the fraction of the enzyme in the input state is lower in preparations expressed at 25°C than in preparations expressed at 22 or 23°C. In some preparations the fractional population of the input state is over 60% even 10 minutes after dilution into buffer. If eNOS is expressed at 25°C, little or no input state can be observed. It is easy for eNOS to lose the ability to form the input state, and such preparations are inactive. Preparations with significant input state populations respond to calmodulin in the same way (shift towards open conformations). Figure 5 illustrates these effects. Fig. 5A compares binding of immobilized calmodulin to several eNOS preparations at 10 nM as previously described [40]; the best preparations bind with high apparent affinity and give large shifts comparable to the top trace shown. Reduction in CaM binding without loss of heme or flavin can be observed after exposure to mild alkaline conditions (pH = 8.8) or after incubation at moderate temperature. The fluorescence decay of a high quality eNOS preparation was shown in Figure 2, and the fractional occupation of each state (amplitude) is given in Table 1. Figure 5B shows the fractional occupations of states in a partially CaM responsive preparation comparable to the one in the middle trace. We cannot conduct binding and fluorescence experiments under identical conditions, but it is clear that eNOS preparations that bind CaM poorly are mainly in open states, and that the individual domains retain their prosthetic groups. Preparations that are particulate respond poorly to CaM but often have large input state components; this can happen if the concentration of glycerol falls below 5%.

**Figure 5.**
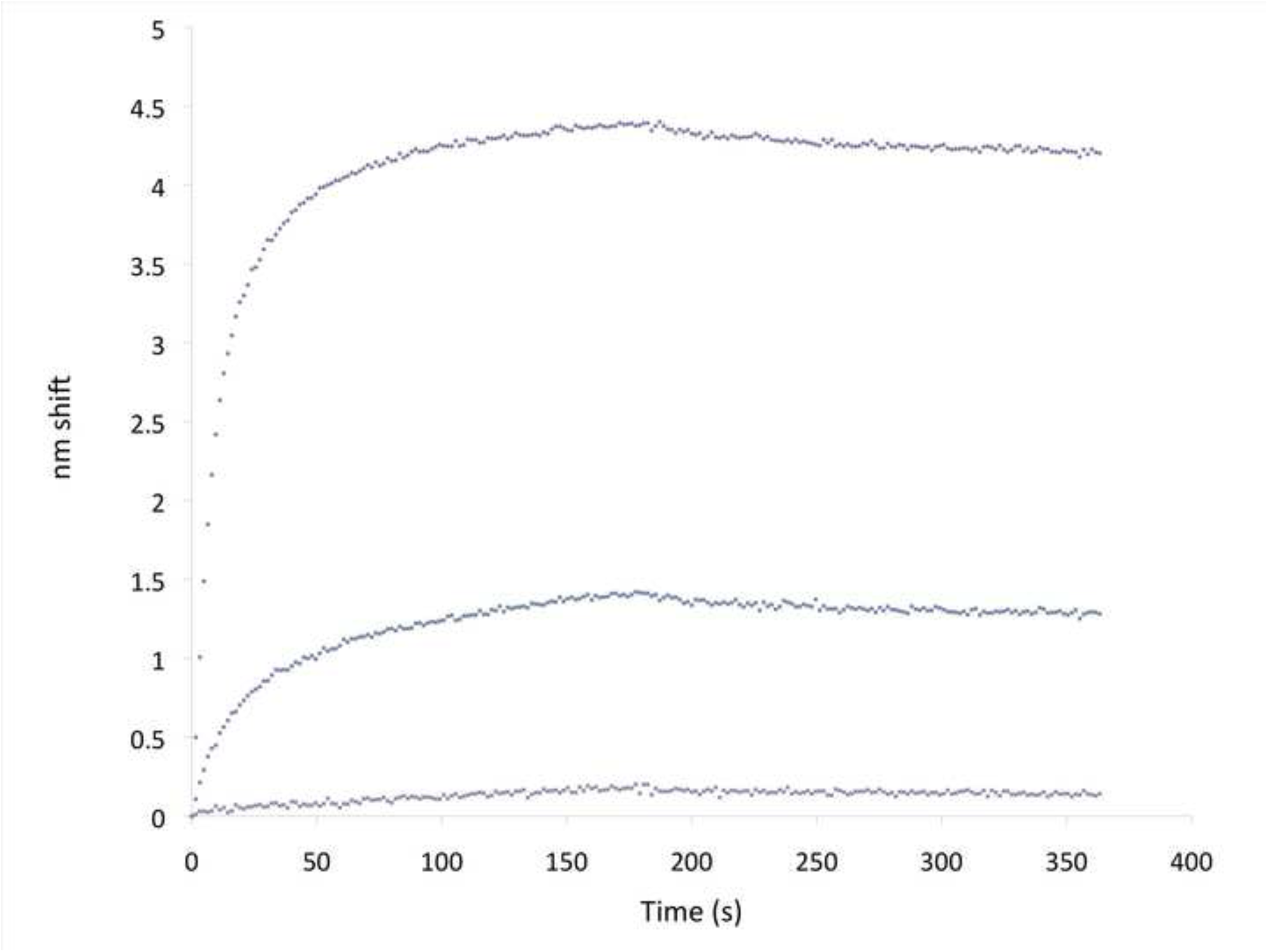
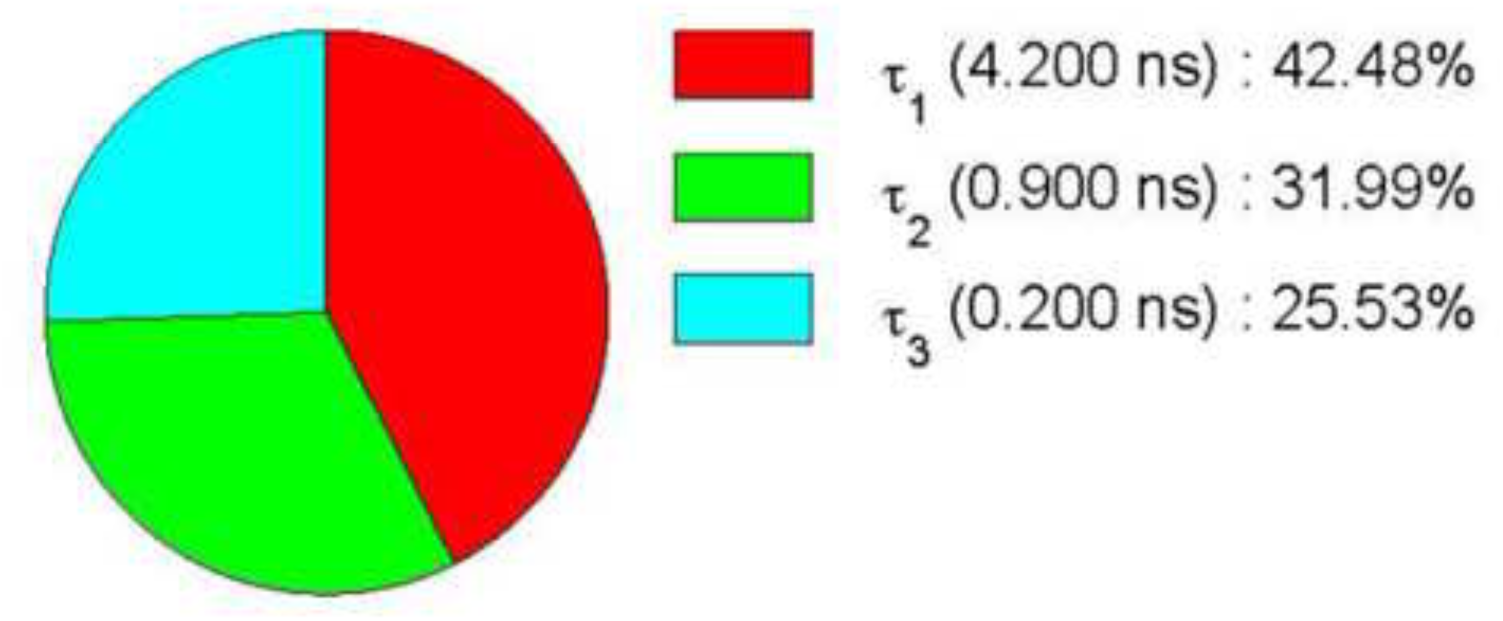
(A) Biolayer interferometry traces showing binding of three different eNOS preparations to immobilized CaM. The top trace is a fresh preparation with high activity. The middle trace is a preparation after repeated freezing and thawing. The lowest trace was exposed to high pH. The association phase was 0-180 s and was followed by 180 s of dissociation. (B) Fractional decay amplitudes of FMN fluorescence decay components of an eNOS preparation comparable to one in the center trace in (A).

As pointed out by Craig et al [54] in the course of kinetics investigations that yielded significant insights, the presence of NADP^+^ has a major effect on the conformational flexibility of NOS reductase units, imposing a ‘conformational lock’. Reduction of cytochrome c by NOS reductase domains is orders of magnitude slower in the presence of NADP^+^/NADPH than in the absence of pyridine nucleotides (by dithionite reduced reductase), but this inhibition is released by Ca^2+^/CaM.

Figure 6 shows the effects of NADP^+^ and NADPH on the fluorescence decay of FMN in nNOS holoenzyme. Addition of NADP^+^ in the absence of CaM causes a modest increase in the fraction of the enzyme in the short lifetime input state. Addition of NADP^+^ to CaM activated eNOS and nNOS also shifts the population towards the short lifetime input state. Ca^2+^/CaM addition in the presence of NADP^+^ can produce very different effects than Ca^2+^/CaM addition to NADP^+^ free enzyme. In some experiments, Ca^2+^/CaM enhances the effect of NADP^+^ on the conformational distribution, further increasing the input state fraction. This apparent paradoxical effect is related to the lack of EDTA reversibility of CaM effects: the principal effect of CaM activation is not to favor one conformation, but to enable conformational transitions. NADP^+^ appears to favor the input state, and CaM activation removes kinetic barriers to allow a larger fraction of the enzyme to reach this state. CaM alone destabilizes the input state, and NADP added after CaM partially reverses this effect. The interactions of CaM and pyridine nucleotide binding are complex.

**Figure 6.**
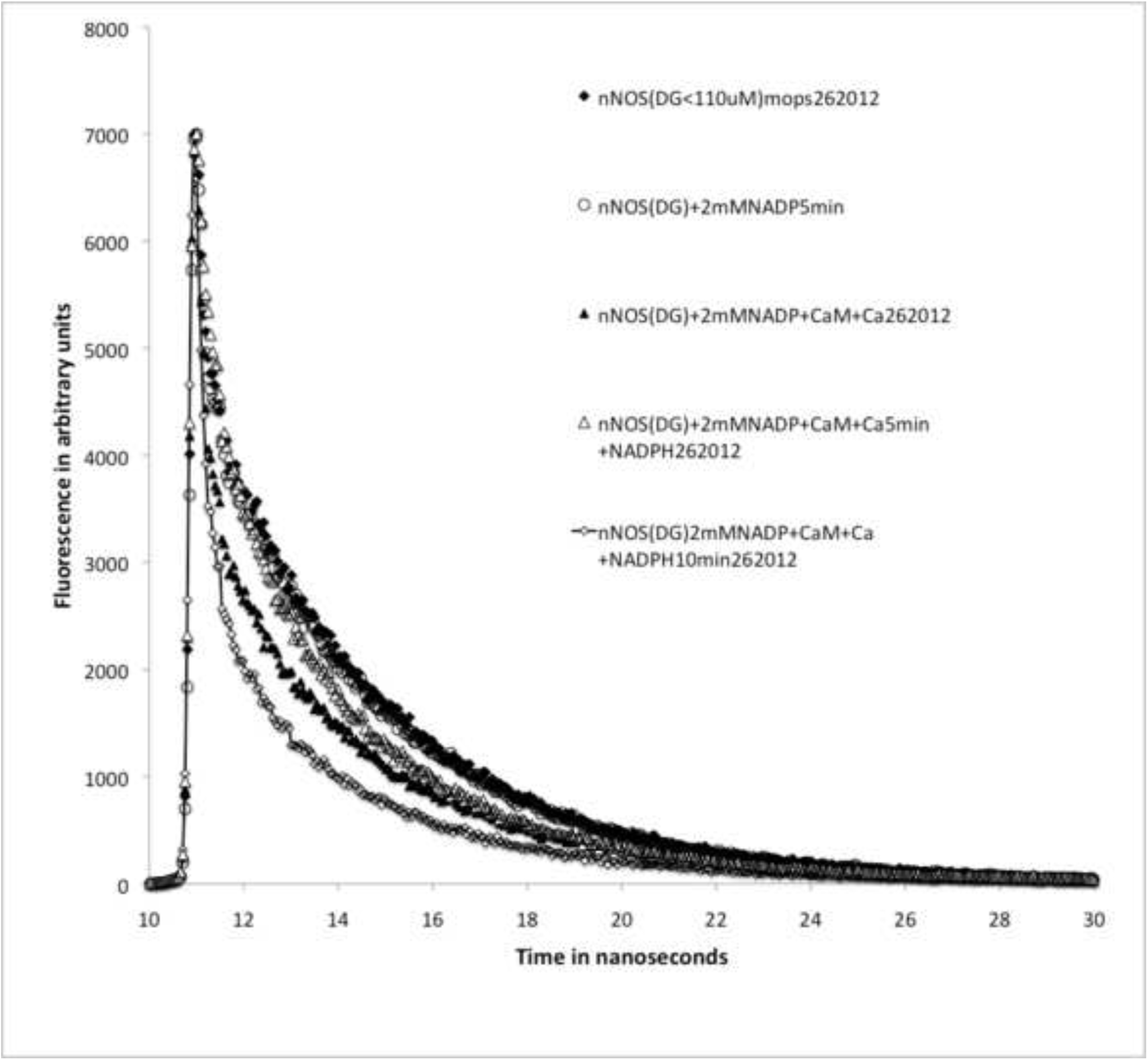
Effects of NADP^+^ and NADPH and CaM on the fluorescence decay of FMN in nNOS holoenzyme, and the paradoxical effect of CaM additional after NADP^+^. ♦, 2 µM nNOS; Ο, + 2mM NADP; ▴, + 2 mM NADP + 4 µM CaM + 10 µM Ca^2+^; ?,+ 2mM NADP + 4 µM CaM + 10 µM Ca^2+^ + 0.2mM NADPH; ◊, + 2mM NADP + 4 µM CaM + 10 µM Ca^2+^ + 0.2 mM NADPH after 10 minutes.

Addition of NADPH in a molar ratio of 1:10 with NADP^+^ produces a very sharp decay composed primarily of short lifetime states; the instantaneous fluorescence (the fluorescence intensity during the duration of the laser pulse) is typically three or four times lower than in oxidized samples, and addition of NADPH alone results in very low levels of flavin fluorescence because the reduced flavins do not absorb in the excitation regions used. The results in samples poised with NADP^+^ and NADPH cannot be interpreted in terms of conformational equilibria alone because the enzyme has a large number of potential redox states as well as a complex conformational manifold. As shown in Figure 7, neither NADPH nor NADP^+^ has a significant effect on the binding of CaM to eNOS, and no significant effects were observed in parallel experiments with nNOS.

**Figure 7.**
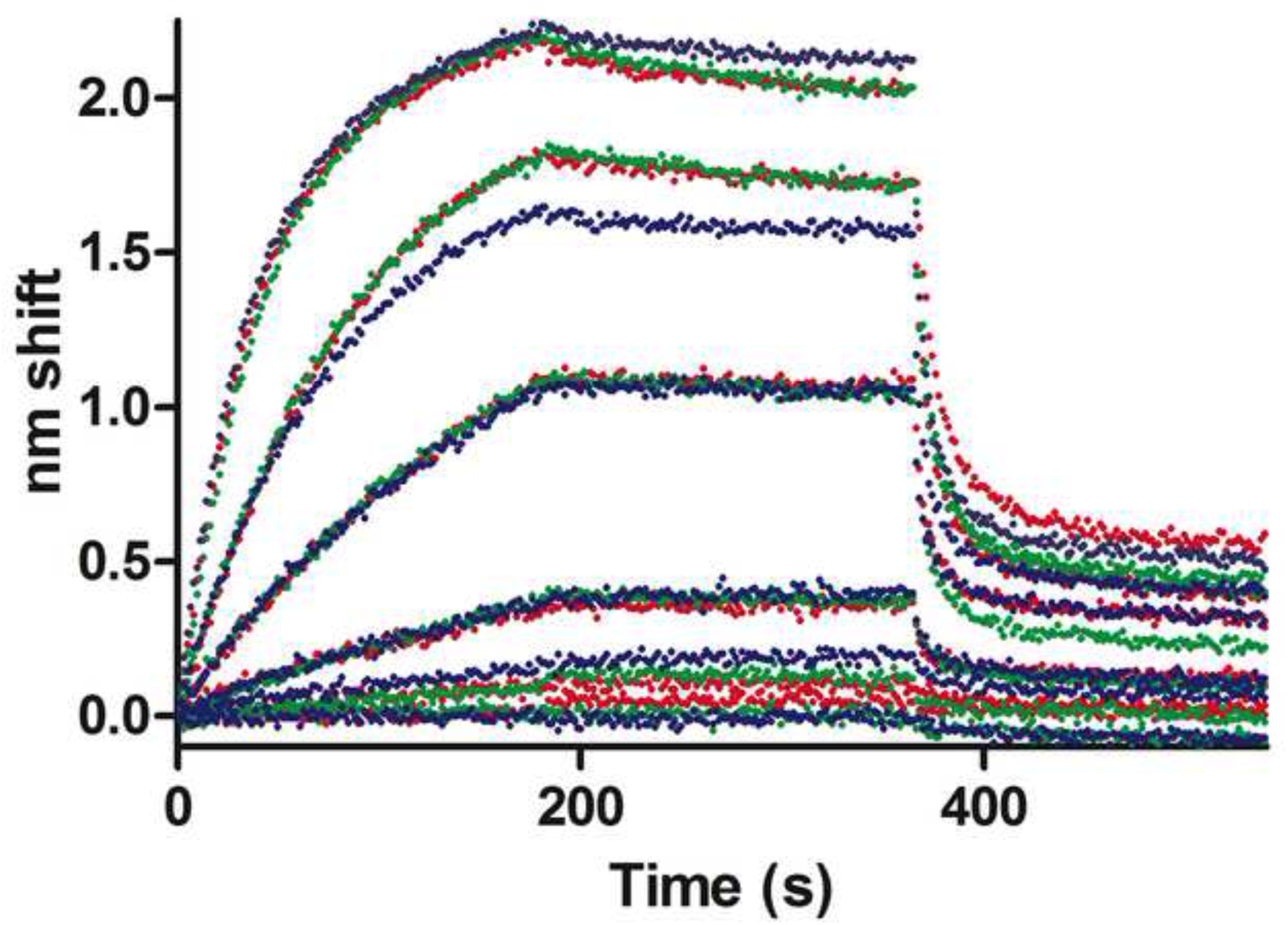
Biolayer interferometry traces showing that NADP^+^ and NADPH do not affect CaM binding to eNOS. Red, no addition; blue, 1 mM NADP; green, 1 mM NADPH. From the top, eNOS concentrations were 150 nM, 75 nM, 38 nM, 19 nM, and 0 nM. The association phase was 0-180s, followed by 180 s of dissociation. Sensors were then dipped into buffer containing 10 mM EDTA to observe rapid dissociation (~360-440 s).

## Discussion

Calmodulin activation of nNOS and eNOS results in multiple effects that are manifested in increased activities for NO synthesis and cytochrome c reduction (a proxy for electron transfer in the reductase unit), and increased FMN/heme electron transfer in oxyFMN. Here we see that these activity changes are the result of changes in the conformational manifold of the enzyme; we observe change in the distribution of states characterized by very different FMN environments. Strictly speaking, these changes are secondary to changes in the rates of interconversion of the conformational states; the equilibrium constants are the ratios of forward and reverse rate constants.

An important factor that governs the time frame of kinetic changes is the presence of a large number of open states, many of which are not in rapid equilibrium with the input and output states, and can be viewed as being unproductive ‘side paths’ in the conformational/catalytic cycle. Most of the increase in fluorescence caused by CaM/Ca^2+^ addition can be fit reasonably well by an exponential with a half time of 30 s, although there are minor slow components. On the other hand, activation of electron transfer to oxygenase heme by stopped flow mixing with concentrated CaM takes no more than 20 ms, consistent with diffusion limited CaM binding.

Clearly, activation is not the result of the increased population of open states, which dominate steady state fluorescence increases. Instead, activation corresponds to switching on conformational transitions that populate the conformational manifold, producing a ‘cloud’ of open states, many of which do not contribute to the catalytic cycle. Because open states, with their long lifetimes, have a much higher quantum yield than the majority input state, an extra 10-15% of total enzyme in these states causes a large increase in total fluorescence. Activation switches on the conformational processes that populate these states over seconds and minutes. The output state is reached in ~ 50 ms, based on stopped flow results.

As in iNOS, the low fluorescence intensities of eNOS and nNOS holoenzymes are the result of FMN-FAD input state pairs in the majority conformational state, which likely closely corresponds to the crystal structures of nNOS reductase domains and P450 reductase. Most of the observed FMN fluorescence is due to other conformational states, and because the high fluorescence yield states are minority states the holoenzymes have relatively weak steady state fluorescence. In contrast, the FMN fluorescence of oxyFMN constructs that lack the FAD and NADPH binding domains is strong because they cannot form the highly quenched input state.

The FMN fluorescence lifetime states of eNOS and nNOS closely correspond to those we recently reported for iNOS constructs [36], and the assignment of the shortest lifetime (90 ps) state to the input state, the 0.9 ns state to the output state, and the longer lifetime states to open conformation appears to hold up well. A major difference between eNOS and nNOS on one hand and iNOS on the other is that in iNOS holoenzyme the level of output state is too low to be reliably detected, and must be less than 5%. In eNOS and nNOS holoenzymes in the presence of Ca^2+^ and CaM the output state is readily detected and accounts for 12-15% of the total population. The iNOS output state can only be detected in constructs (e.g., oxyFMN) designed to suppress the input state.

This suggests that FMN/heme electron transfer in iNOS occurs in a state that is a small minority (< 5%) of the total conformational manifold. In eNOS and nNOS the control elements (autoinhibitory insertion and C terminal restrictor) lock the reductase unit into the input state, in part through a network of H bonds that can be visualized using crystallographic results. Why then does iNOS, lacking an AI and with a shorter C terminal element, have a smaller output state population than nNOS and eNOS?

A solution to this apparent paradox can be inferred by considering the obligatory conformational cycle associated with reductase catalysis. In iNOS and in CaM-activated eNOS and nNOS, cytochrome c reduction is orders of magnitude faster than oxygenase heme reduction; release of the FMN binding domain is not rate limiting for catalytic delivery of electrons to the oxygenase active site. In oxyFMN constructs, FMN-heme electron transfer occurs in a single kinetic phase that is much faster than heme reduction in holoenzyme, so association of open conformation FMN binding domains with oxygenase domains does not appear to be rate limiting *per se*.

As pointed out in our iNOS study [36], the slowest step is instead passage of the FMN binding domain through a conformational bottleneck that connects the open states in rapid equilibrium with the input states with other open states in rapid equilibrium with output states. CaM binding appears to affect all these processes, in part through steric interactions that exclude unproductive states. The ‘bottleneck’ description is meant to indicate the slow step in the kinetic mechanism, but does not imply that the open states in the bottleneck between the input and output states are necessarily higher in energy than the states adjacent to the input and output states. As shown in Figure 8, we envision the input and output states as deep, narrow valleys in the conformational landscape, while there are many open states. Many open states represent orientations of the FMN binding domain that are not on the pathway between electron transfer active states, and the kinetic bottleneck may in part be the consequence the relatively small number of ‘productive’ open conformations. The long timeframe of development of steady state fluorescence after CaM binding and activation is a consequence of the existence of open states far from productive pathways; a minute after CaM activation, a sizeable fraction (likely at least 20%) of the enzyme population is in such states, and because of their high fluorescence yield (about 50 times that of the input state, and five times that of the minority output state) these states dominate the steady state fluorescence. The full development of steady state fluorescence after activation is not a report of activation but rather an eventual consequence of it.

**Figure 8.**
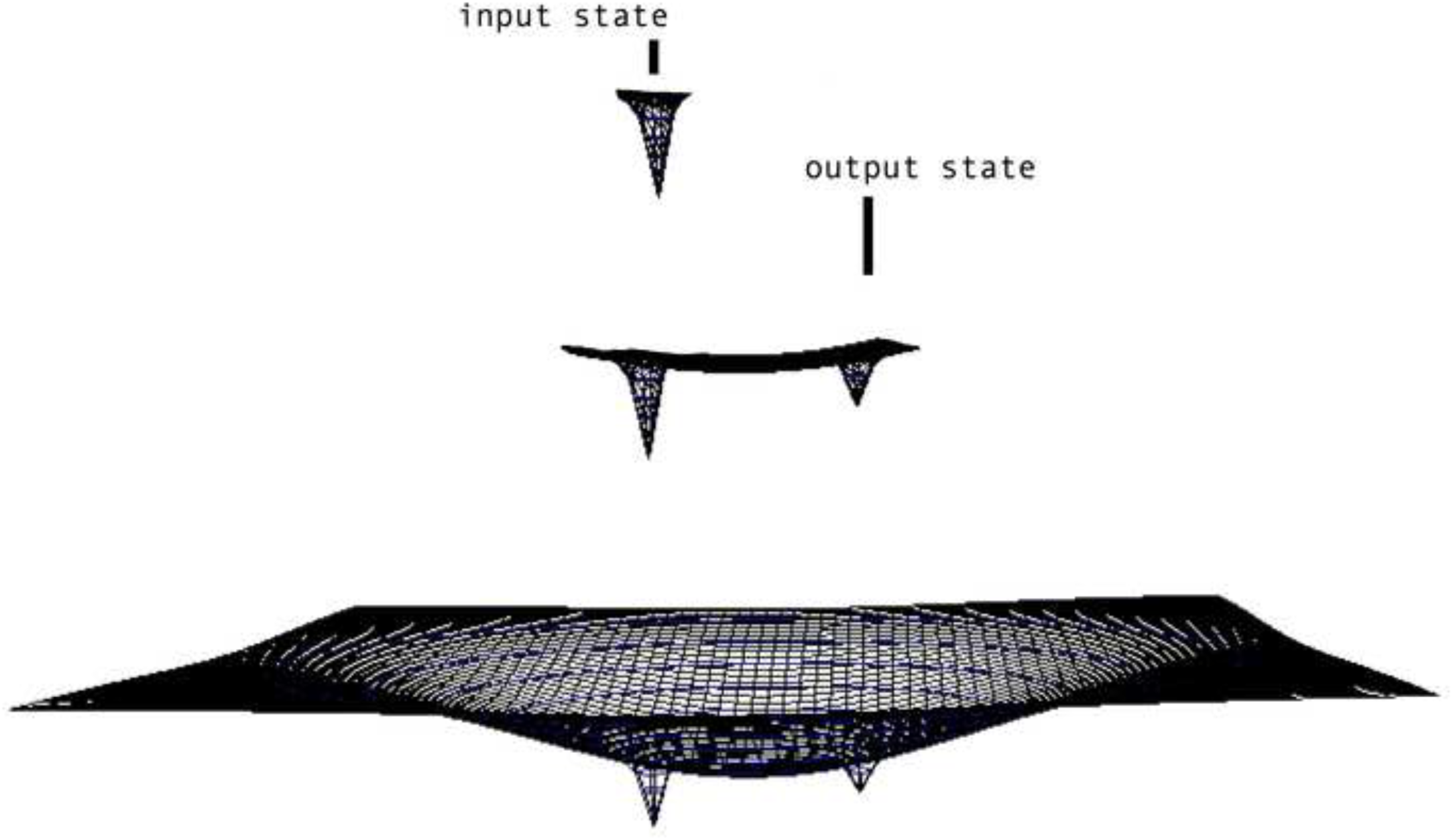
Schematic of conformational landscape for NOS reductase, showing sharp input and output state wells and broad shallow open state bowl. Prior to activation, most NOS molecules are in the input state (top), which has very weak steady state fluorescence. Activation provides rapid access to open configurations near the input state (~10 ms), and the output state is reached within ~50 ms (middle). Tethered diffusion of the FMN binding domain is not limited to open configurations near the input and output states, and more distant configurations are accessed on the time scale of seconds and minutes. Although at these configurations are attained by a minority of the enzymes, they dominate the steady state fluorescence because of their long lifetimes.

During turnover, the rate of electron delivery is not solely determined by the concentration of the output state in the FMNH_2_-heme (FeIII) redox state, but rather is determined by King-Altman steady state considerations that include redox processes and by the return of the FMN binding domain to the input state. Considering the conformational equilibrium of the oxidized enzyme, it appears that control elements not only stabilize the input state (this is clearly true from kinetic results), but also interfere with the return of the oxygenase domain associated FMN binding domain to the input state configuration. A glance at the crystal structure of the input state of the nNOS reductase unit shows why these elements might slow down association as well as release; the C terminal element guards one edge of the binding site for the FMN binding domain, and the AI entangles the other edge, complicating the interactions for successful docking. NOS control elements appear to inhibit the formation of the input state as well as its breakup. This leaves a larger fraction of the enzyme in the output state than in iNOS.

The interconversion of conformational states is slow on the time scale of fluorescence lifetime experiments (about 4 ns because we can resolve the discrete fluorescence lifetime states and the corresponding conformational states). Transitions from open conformations to short lifetime states must occur on time scales slower than ~ 8 ns; this might contribute to microheterogeneity of the long lifetime states, since open states adjacent to the short lifetime states might exchange with them rapidly enough to contribute to relaxation. As pointed out in our paper on iNOS [36], in kinetic experiments the rate of interconversion must be fast enough to average kinetic components of electron transfer. Interconversion must occur in the time regime of 10^-3^ to 10^-8^ seconds because it is faster than electron transfer and slower than fluorescence decay.

As stated earlier, NADP^+^/NADPH binding has a significant effect on the conformational repertoire of the reductase domains in the absence of CaM [54]. In the absence of NADPH, dithionite reduced, CaM-free NOS can rapidly reduce cytochrome c (25° C rate constant ~ 15 M^-^ 1 s^-1^), but NADPH reduced NOS in the absence of CaM reduces cytochrome c much more slowly (25° C rate constant ~ 1.1 M^-1^ s^-1^). In the presence of CaM, both rates are just over 35 M^-1^ s^-1^. This was attributed to a ‘conformational lock’ imposed by tightly bound NADPH. However, a rapid minor phase of cytochrome c reduction in NADPH reduced NOS without bound CaM was noted. This corresponded to much less than 1 electron per NOS, but appears consistent with 10-20% of the enzyme donating an electron on the timescale of 10 - 20 ms, more than an order of magnitude faster than the large slow component. This is likely the result of a significant fraction of the enzyme locked into an open state that slowly equilibrates with the input state in the absence of CaM.

Kinetic considerations and physical measurements suggest a heme-FMN edge to distance of 13.5 ± 1.5 Å in the iNOS output state [24, 55] and this is consistent with previous experiments using nNOS oxyFMN constructs [23, 37]. The output states of iNOS, eNOS and nNOS, having similar fluorescence lifetimes as a result of heme quenching, are likely to be very similar [36, 39].

The relatively slow electron transfer rate in eNOS is not the result of an output state very different from the much more rapid electron transfer systems in nNOS and iNOS but to a King-Altman steady state in which a much smaller portion of the enzyme is in the kinetically competent state for heme reduction during turnover. This is likely to be due largely to the eNOS control elements, but contributions from interactions between domain edges and differences in linker regions may also contribute.

We have demonstrated that control of the nitric oxide synthase signal generators by Ca^2+^/CaM involves the control of major conformational reorientation of the FMN binding domain by CaM binding. The similarity of the FMN fluorescence lifetimes in the corresponding conformational states of all three isoforms is strong evidence that these states are similar, and that the different kinetic properties of these isoforms is due to differences in the rates of interconversion of these states, which results in the changes in steady state and equilibrium distributions in the conformational manifold.

## Acknowledgements

John C. Salerno passed away on Dec. 25, 2015. As this is likely his final work, his coauthors wish to express our gratitude for John’s invaluable contributions not only to science, but to our lives and careers as collaborator, mentor and friend. We gratefully acknowledge access to the facilities at the Center for Fluorescence Spectroscopy at University of Maryland School of Medicine in obtaining some of the fluorescence lifetime measurements. We thank Thomas Poulos and Huiying Li of University of California-Irvine for provision of samples and helpful comments on the manuscript. We also thank Cold Spring Harbor for creating and maintaining BioRxiv and thus providing a repository for this work.

## References

1. Furchgott RF. Vascular smooth muscle, peptides, autonomic nerves and endothelium. In: Vanhoutte PM, editor. Vasodilation. New York: Raven Press; 1988. p. 401–4.

2. Ignarro LJ, Buga GM, Wood KS, Byrns RE, Chaudhuri G. Endothelium-derived relaxing factor produced and released from artery and vein is nitric oxide. Proc Natl Acad Sci USA. 1987;84(9265–9269).

3. Garthwaite J, Charles SL, Chess-Williams R. Endothelium-derived relaxing factor release on activation of NMDA receptors suggests role as intercellular messenger in the brain. Nature (London). 1988;336:385–8.

4. Gally JA, Montague PR, Reeke JGN, Edelman GM. Nitric oxide: linking space and time in the brain. Proc Natl Acad Sci USA. 1990;87(3547–3551).

5. Bredt DS, Hwang PM, Snyder SH. Localization of nitric oxide synthase indicating a neural role for nitric oxide. Nature. 1990;347(768–770).

6. Abu-Soud HM, Stuehr DJ. Nitric oxide synthases reveal a role for calmodulin in controlling electron transfer. Proc Natl Acad Sci USA. 1993;90(10769–10772).

7. Ghosh DK, Salerno JC. Nitric oxide synthases: domain structure and alignment in enzyme function and control. Front Biosci. 2003;8:193–209.

8. Daff S. NO synthase: structures and mechanisms. Nitric Oxide. 2010;23:1–11.

9. Li H, Poulos TL. Structure-function studies on nitric oxide synthases. J Inorg Biochem. 2005;99:293–305.

10. Salerno JC, Ghosh DK. Space, time and nitric oxide -- neuronal nitric oxide synthase generate signal pulses. FEBS J. 2009;276:6677–88.

11. Roman LJ, Martasek P, Masters BS. Intrinsic and extrinsic modulation of nitric oxide synthase activity. Chem Rev. 2002;102:1179–90.

12. Stuehr DJ, Tejero J, Haque MM. Structural and mechanistic aspects of flavoproteins: electron transfer through the nitric oxide synthase flavoprotein domain. FEBS J. 2009;276(15):3959–74.

13. Campbell MG, Smith BC, Potter CS, Carrager B, Marletta MA. Molecular architecture of mammalian nitric oxide synthases. Proc Natl Acad Sci USA. 2014;111:E3614-623.

14. Smith BC, Underbakke ES, Kulp DW, Schief WR, Marketta MA. Nitric oxide synthase domain interfaces regulate electron transfer and calmodulin activation. Proc Natl Acad Sci USA. 2013;110:E3577–86.

15. Salerno JC, Harris DE, Irizarry K, Patel B, Morales AJ, Smith SM, et al. An autoinhibitory control element defines calcium-regulated isoforms of nitric oxide synthase. J Biol Chem. 1997;272:29769–77.

16. Daff S. Calmodulin-dependent regulation of mammalian nitric oxide synthase. Biochem Soc Trans. 2003;31:502–5.

17. Tiso M, Tejero J, Panda K, Aulak KS, Stuehr DJ. Versatile regulation of neuronal nitric oxide synthase by specific regions of its C-terminal tail. Biochemistry. 2007;46(50):14418–28.

18. Roman LJ, Martasek P, Miller RT, Harris DE, de la Garza MA, Shea TM, et al. The C termini of constitutive nitric-oxide synthases control electron flow through the flavin and heme domains and affect modulation by calmodulin. J Biol Chem. 2000;275(38):29225–32.

19. Knowles RG, Merrett M, Salter M, Moncada S. Differential induction of brain, lung and liver nitric oxide synthase by endotoxin in the rat. Biochem J. 1990;270(3):833–6.

20. Marletta MA, Yoon PS, Iyengar R, Leaf CD, Wishnok JS. Macrophage oxidation of L-arginine to nitrite and nitrate: nitric oxide is an intermediate. Biochemistry. 1988;27(24):8706–11.

21. Stuehr DJ, Gross SS, Sakuma I, Levi R, Nathan CF. Activated murine macrophages secrete a metabolite of arginine with the bioactivity of endothelium-derived relaxing factor and the chemical reactivity of nitric oxide. J Exp Med. 1989;169(3):1011–20.

22. Cho HJ, Xie QW, Calaycay J, Mumford RA, Swiderek KM, Lee TD, et al. Calmodulin is a subunit of nitric oxide synthase from macrophages. J Exp Med. 1992;176(2):599–604.

23. Ghosh DK, Holliday MA, Thomas C, Weinberg JB, Smith SM, Salerno JC. Nitric-oxide synthase output state. Design and properties of nitric-oxide synthase oxygenase/FMN domain constructs. J Biol Chem. 2006;281(20):14173–83.

24. Feng C, Thomas C, Holliday MA, Tollin G, Salerno JC, Ghosh DK, et al. Direct measurement by laser flash photolysis of intramolecular electron transfer in a two-domain construct of murine inducible nitric oxide synthase. J Am Chem Soc. 2006;128(11):3808–11.

25. Feng C, Tollin G, Hazzard JT, Nahm NJ, Guillemette JG, Salerno JC, et al. Direct measurement by laser flash photolysis of intraprotein electron transfer in a rat neuronal nitric oxide synthase. J Am Chem Soc. 2007;129(17):5621–9.

26. Li H, Das A, Sibhatu H, Jamal J, Sligar SG, Poulos TL. Exploring the electron transfer properties of neuronal nitric-oxide synthase by reversal of the FMN redox potential. J Biol Chem. 2008;283(50):34762–72.

27. Ilagan RP, Tejero J, Aulak KS, Ray SS, Hemann C, Wang ZQ, et al. Regulation of FMN subdomain interactions and function in neuronal nitric oxide synthase. Biochemistry. 2009;48(18):3864–76.

28. Yokom AL, Miorishima Y, Lau M, Su M, Glukhova A, Osawa Y, et al. Architecture of the nitric-oxide synthase holoenzyme reveals large conformational changes and a calmodulin-driven release of the FMN domain. J Biol Chem. 2014;289:16855–65.

29. Volkmann N, Martasek P, Roman LJ, Xu XP, Page C, Swift M, et al. Holoenzyme structures of endothelial nitric oxide synthase - an allosteric role for calmodulin in pivoting the FMN domain for electron transfer. J Struct Biol. 2014;188:46–54.

30. Sheng Y, Zhong L, Guo D, Lau G, Feng C. Insight into structural rearrangements and interdomain interactions related to electron transfer between flavin mononucleotide and heme in nitric oxide synthase: A molecular dynamics study. J Inorg Biochem. 2015;153:186–96.

31. Hollingsworth SA, Holden JK, Poulos TL. Elucidating nitric oxide synthase domain interactions by molecular dynamics. Protein Sci. 2016;25(2):374–82.

32. Jáchymová M, Martásek P, Panda S, Roman LJ, Panda M, Shea TM, et al. Recruitment of governing elements for electron transfer in the nitric oxide synthase family. Proc Natl Acad Sci USA. 2005;102(44):15833–8.

33. Gachhui R, Presta A, Bentley DF, Abu-Soud HM, McArthur R, Brudvig G, et al. Characterization of the reductase domain of rat neuronal nitric oxide synthase generated in the methylotrophic yeast Pichia pastoris. Calmodulin response is complete within the reductase domain itself. J Biol Chem. 1996;271(34):20594–602.

34. Newman E, Spratt DE, Mosher J, Cheyne B, Montgomery HJ, Wilson DL, et al. Differential activation of nitric-oxide synthase isozymes by calmodulin-troponin C chimeras. J Biol Chem. 2004;279(32):33547–57.

35. Roman LJ, Masters BS. Electron transfer by neuronal nitric oxide synthase is regulated by concerted interaction of calmodulin and two intrinsic regulatory elements. J Biol Chem. 2006;281(32):23111–8.

36. Ghosh DK, Ray K, Rogers AJ, Nahm NJ, Salerno JC. FMN fluorescence in inducible NOS constructs reveals a series of conformational states involved in the reductase catalytic cycle. FEBS J. 2012;279(7):1306–17.

37. Feng C, Tollin G, Holliday MA, Thomas C, Salerno JC, Enemark JH, et al. Intraprotein electron transfer in a two-domain construct of neuronal nitric oxide synthase: the output state in nitric oxide formation. Biochemistry. 2006;45(20):6354–62.

38. Li W, Fan W, Elmore BO, Feng C. Effect of solution viscosity on intraprotein electron transfer between the FMN and heme domains in inducible nitric oxide synthase. FEBS Lett. 2011;585(16):2622–6.

39. Salerno JC, Ray K, Poulos TL, Li H, Ghosh DK. Calmodulin activates neuronal nitric oxide synthase by enabling transitions between conformational states. FEBS Lett. 2013;587(1):44–7.

40. Bredt DS, Hwang PM, Glatt CE, Lowenstein C, Reed RR, Snyder SH. Cloned and expressed nitric oxide synthase structurally resembles cytochrome P-450 reductase. Nature. 1991;351(6329):714–8.

41. Roman LJ, Sheta EA, Martásek P, Gross SS, Liu Q, Masters BS. High-level expression of functional rat neuronal nitric oxide synthase in Escherichia coli. J Biol Chem. 1995;92(18):8428–32.

42. Ghosh DK, Rashid MB, Crane B, Taskar V, Mast M, Misukonis MA, et al. Characterization of key residues in the subdomain encoded by exons 8 and 9 of human inducible nitric oxide synthase: a critical role for Asp-280 in substrate binding and subunit interactions. Proc Natl Acad Sci USA. 2001;98(18):10392–7.

43. Gross SS. Microtiter plate assay for determining kinetics of nitric oxide synthesis. Methods Enzymol. 1996;268:159–68.

44. Martásek P, Liu Q, Liu J, Roman LJ, Gross SS, Sessa WC, et al. Characterization of bovine endothelial nitric oxide synthase expressed in E. coli. Biochem Biophys Res Comm. 1996;219(2):359–65.

45. Ray K, Szmacinski H, Lakowicz JR. Enhanced fluorescence of proteins and label-free bioassays using aluminum nanostructures. Anal Chem. 2009;81(15):6049–54.

46. Lakowicz JR. Principles of Fluorescence Spectroscopy. 3rd edition ed. New York: Springer; 2006.

47. McMurry JL, Chrestensen CA, Scott IM, Lee EW, Rahn AM, Johansen AM, et al. Rate, affinity and calcium dependence of CaM binding to eNOS and nNOS: effects of phosphorylation. FEBS J. 2011;278(24):4943–54.

48. Narayanasami R, Nishimura JS, McMillan K, Roman LJ, Shea TM, Robida AM, et al. The influence of chaotropic reagents on neuronal nitric oxide synthase and its flavoprotein module. Urea and guanidine hydrochloride stimulate NADPH-cytochrome c reductase activity of both proteins. Nitric Oxide. 1997;1(1):39–49.

49. Brunner K, Tortschanoff A, Hemmens B, Andrew PJ, Mayer B, Kungl AJ. Sensitivity of flavin fluorescence dynamics in neuronal nitric oxide synthase to cofactor-induced conformational changes and dimerization. Biochemistry. 1998;37(50):17545–53.

50. Bastiaens PI, Bonants PJ, Müller F, Visser AJ. Time-resolved fluorescence spectroscopy of NADPH-cytochrome P-450 reductase: demonstration of energy transfer between the two prosthetic groups. Biochemistry. 1989;28(21):8416–25.

51. Climent T, González-Luque R, Merchán M, Serrano-Andrés L. Theoretical insight into the spectroscopy and photochemistry of isoalloxazine, the flavin core ring. J Phys Chem A. 2006;110(50):13584–90.

52. Wang M, Roberts DL, Paschke R, Shea TM, Masters BS, Kim JJ. Three-dimensional structure of NADPH-cytochrome P450 reductase: prototype for FMN- and FAD-containing enzymes. Proc Natl Acad Sci USA. 1997;94(16):8411–6.

53. Garcin ED, Bruns CM, Lloyd SJ, Hosfield DJ, Tiso M, Gachhui R, et al. Structural basis for isozyme-specific regulation of electron transfer in nitric-oxide synthase. J Biol Chem. 2004;279(36):37918–27.

54. Craig DH, Chapman SK, Daff S. Calmodulin activates electron transfer through neuronal nitric-oxide synthase reductase domain by releasing an NADPH-dependent conformational lock. J Biol Chem. 2002;277(37):33987–4.

55. Astashkin AV, Elmore BO, Fan W, Guillemette JG, Feng C. Pulsed EPR determination of the distance between heme iron and FMN centers in a human inducible nitric oxide synthase. J Am Chem Soc. 2010;132(34):12059–67.

